# Zebrafish spinal cord oligodendrocyte formation requires *boc* function

**DOI:** 10.1101/2021.01.31.429029

**Authors:** Christina A. Kearns, Macie Walker, Andrew M. Ravanelli, Kayt Scott, Madeline R. Arzbecker, Bruce Appel

## Abstract

The axis of the vertebrate neural tube is patterned, in part, by a ventral to dorsal gradient of Shh signaling. In the ventral spinal cord, Shh induces concentration-dependent expression of transcription factors, subdividing neural progenitors into distinct domains that subsequently produce distinct neuronal and glial subtypes. In particular, progenitors of the pMN domain express the bHLH transcription factor Olig2 and produce motor neurons followed by oligodendrocytes, the myelinating glial cell type of the central nervous system. In addition to its role in patterning ventral progenitors, Shh signaling must be maintained through development to specify pMN progenitors for oligodendrocyte fate. Using a forward genetic screen in zebrafish for mutations that disrupt development of oligodendrocytes, we identified a new mutant allele of *boc*, which encodes a type I transmembrane protein that functions as a coreceptor for Shh. Embryos homozygous for the *boc*^*co25*^ allele, which creates a missense mutation in a Fibronectin type III domain that binds Shh, have normally patterned spinal cords but fail to maintain pMN progenitors, resulting in a deficit of oligodendrocytes. Using a sensitive fluorescent detection method for in situ RNA hybridization, we found that spinal cord cells express *boc* in a graded fashion that is inverse to the gradient of Shh signaling activity and that *boc* function is necessary to maintain pMN progenitors by shaping the Shh signaling gradient.

## Introduction

In the central nervous system (CNS) of vertebrate animals, the speed and timing of axon electrical impulses and neuron health are supported by myelin, a modified membrane produced by glial oligodendrocytes (Simons and Nave 2016). In the spinal cord, most oligodendrocytes arise from ventrally positioned progenitor cells (Warf *et al.* 1991; Noll and Miller 1993). These progenitors, known as pMN cells, express the basic helix loop helix (bHLH) transcription factor Olig2 (Lu *et al.* 2000; Takebayashi *et al.* 2000; Zhou *et al.* 2000; Park *et al.* 2002). pMN progenitors first produce motor neurons and then switch to making oligodendrocytes (Richardson *et al.* 2000; Soula *et al.* 2001). Maintenance of the pMN progenitor population is essential for the formation of appropriate numbers of motor neurons and oligodendrocytes but the mechanistic basis of pMN progenitor maintenance remains unclear.

The notochord, a mesodermal structure ventral to the neural tube, patterns the ventral neural tube by secreting the morphogen Shh (Wijgerde et al., 2002, Echelard et al., 1993; Martí et al., 1995; Roelink et al., 1994). Shh binds the transmembrane receptor Patched (Ptch) thereby relieving Ptch inhibition of Smoothened (Smo) and culminating in changes in transcription of Shh pathway target genes (Briscoe and Thérond 2013). Graded Shh activity induces expression of genes that encode bHLH and homeodomain (HD) proteins at distinct positions on the dorsal-ventral (DV) axis, including Olig2 (Briscoe *et al.* 2000; Dessaud *et al.* 2008, 2010; Kutejova *et al.* 2016). Accordingly, spinal cord cells of mutant mouse embryos lacking Shh failed to express Olig2 (Lu *et al.* 2000; Oh *et al.* 2005). Shh signaling also is required to maintain pMN progenitors through the period of neurogenesis. Chick and zebrafish embryos treated with cyclopamine, which inhibits Shh pathway signaling activity, following neural tube patterning had deficits of Olig2^+^ pMN cells and OPCs (Park *et al.* 2004; Allen *et al.* 2011; Oustah *et al.* 2014; Ravanelli and Appel 2015). Thus, Shh signaling specifies and maintains spinal cord progenitors that give rise to motor neurons and oligodendrocytes.

Importantly, the responsiveness of neural tube cells to Shh is modified by proteins that function as Shh co-receptors, including Gas1, Cdo and Boc. Gas1 is a GPI-anchored cell surface protein with a high affinity for Shh (Lee *et al.* 2001) and Cdo and Boc are Ig super-family members that contain four (Boc) and five (Cdo) extracellular Ig repeats adjacent to fibronectin type III (FNIII) repeats, which are critical for Shh binding (Kang *et al.* 1997, 2002; Tenzen *et al.* 2006; Yao *et al.* 2006). Notochord and floor plate cells transiently express Cdo during early neural tube patterning; subsequently, high levels of Gas1, Cdo and Boc expression are evident within dorsal spinal cord cells (Tenzen *et al.* 2006; Allen *et al.* 2007, 2011). Although mouse embryos carrying inactivating mutations of individual co-receptor genes have rather mild spinal cord patterning abnormalities, double mutant combinations of *Gas1, Cdo* and *Boc* result in a decrease of pMN progenitor populations in late stages of neural development, indicating that pMN progenitor maintenance requires function of these co-receptors (Allen *et al.* 2007, 2011). Furthermore, ectopic expression of these receptors in the chick neural tube caused a Shh-dependent ventralization of the spinal cord, marked by increased dorsal expression of Olig2 and Nkx2.2 (Tenzen *et al.* 2006; Allen *et al.* 2007, 2011). Because co-receptor expression can enhance Shh signaling activity (Tenzen *et al.* 2006; Martinelli and Fan 2007) these observations suggest that Shh co-receptors enhance cell responsiveness to Shh necessary to maintain pMN progenitors.

Here we provide new evidence for the requirement of Boc function in pMN progenitor maintenance and motor neuron and oligodendrocyte formation. From a forward genetic screen in zebrafish we identified a mutant allele of *boc* that caused a deficit of oligodendrocytes. A detailed expression analysis revealed that *olig2*^*+*^ pMN cells co-express *boc* and *ptch2* and that loss of *boc* function alters the gradient of Shh signaling activity. Furthermore, mutant *boc* embryos have deficits of late-born motor neurons and a significant decrease in oligodendrocytes due to a progressive depletion of pMN progenitors. We therefore conclude that Boc function maintains pMN progenitors for motor neuron and oligodendrocyte fate by sustaining appropriate levels of Shh signaling in the pMN domain.

## Results

### Mutation of boc impairs formation of motor neurons and oligodendrocyte lineage cells

In a screen for chemically induced mutations that disrupt oligodendrocyte development we recovered a line, given the allele designation *co25*, for which subsets of larvae produced by sibling incrosses had reduced numbers of dorsally migrated oligodendrocyte lineage cells, marked by *olig2*:EGFP expression, and a variable downward trunk and tail curvature. Whereas less than 25% of larvae produced by pairwise incrosses had trunk curvature, the deficit of oligodendrocyte lineage cells was evident in frequencies approximating a Mendelian ratio (**Figure 1A**). To identify the gene affected by the *co25* mutant allele we performed bulk segregant analysis, which centered the locus at about 50 cM on chromosome 24 of the MGH mapping panel. Located at 44.1 cM on chromosome 24 is *boc,* which encodes an Ig/fibronectin superfamily member that can function as a Shh co-receptor to enhance Shh signaling (Tenzen *et al.* 2006). Because *boc* mutant zebrafish larvae have curved body axes (Bergeron *et al.* 2011) similar to *co25* mutant larvae, we chose *boc* as a candidate gene. Sequencing homozygous mutant embryos revealed a T to G transversion mutation within exon 13 predicted to cause substitution of serine for a conserved isoleucine at amino acid position 721 of the Boc protein (Figure 1B). This substitution lies within the third fibronectin type-III (FNIII(3)) domain of Boc, which can bind to Shh (Tenzen *et al.* 2006). To validate identification of the *co25* allele as a mutation of *boc*, we crossed heterozygous *co25* fish to fish heterozygous for the *boc*^*ty54*^ allele, which introduces a premature stop codon at amino acid position 238 (Bergeron *et al.* 2011). Larval progeny of these crosses had morphological and oligodendrocyte lineage cell phenotypes similar to larvae homozygous for the *co25* allele (Figure 1C). We therefore conclude that the *co25* allele is a loss-of-function mutation of *boc,* which might interfere with the binding interaction of Boc with Shh. A following experiments were performed using the straight tail class of embryos and larvae homozygous for the *boc*^*co25*^ allele.

**Figure 1.**
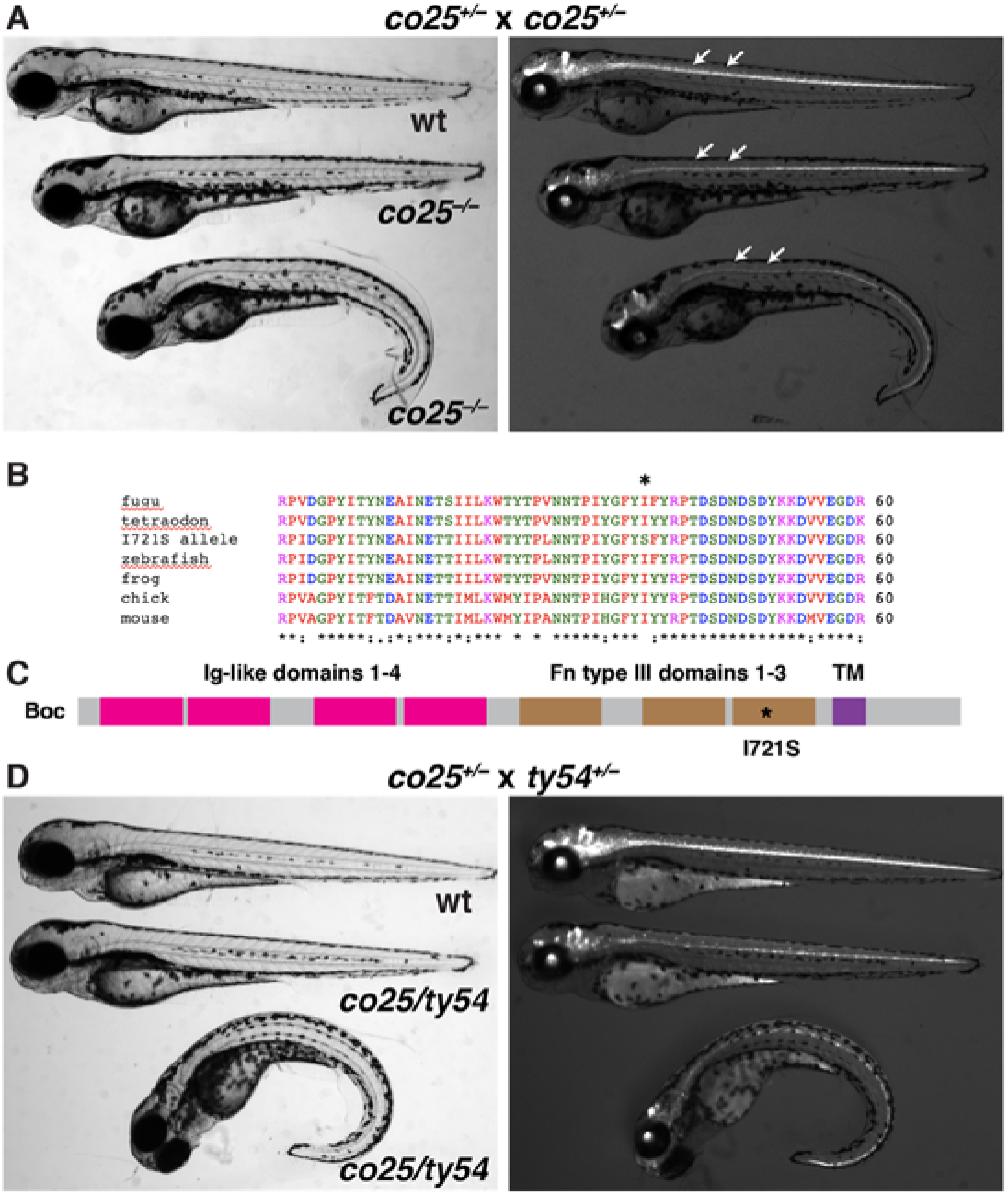
Identification of a new *boc* mutant allele. (A) Brightfield and epifluorescent images of 3 dpf larvae produced by heterozygous *co25* adult zebrafish. Arrows indicate dorsal spinal cord, occupied by OPCs marked by *olig2*:EGFP expression. Straight and curved body homozygous mutant larvae have similar reductions in *olig2*:EGFP fluorescence intensity and OPC number. (B) Amino acid alignment showing the isoleucine to serine substitution at position 721 (asterisk). (C) Domain structure of Boc. The *co25* mutant allele causes substitution of serine for isoleucine within Fn type III domain 3. (D) Complementation test with the *boc*^*ty54*^ allele. Transheterozygous larvae have phenotypes similar to homozygous *co25* mutant larvae.

To confirm that *boc* mutant larvae have fewer oligodendrocyte lineage cells we performed immunohistochemistry and in situ RNA hybridization to detect gene expression that defines the lineage. In the CNS Sox10 expression marks oligodendrocyte lineage cells, consisting of oligodendrocyte precursor cells (OPCs) and oligodendrocytes. Labeling with an antibody specific to Sox10 revealed that *boc* mutant larvae had fewer than 25% of the oligodendrocyte lineage cells characteristic of wild type (Figure 2A-C). Consistent with this result, *boc* mutant larvae had substantially fewer *myrf^+^* pre-myelinating oligodendrocytes and *mbpa^+^* oligodendrocytes than wild-type larvae (Figure 2D-I).

**Figure 2.**
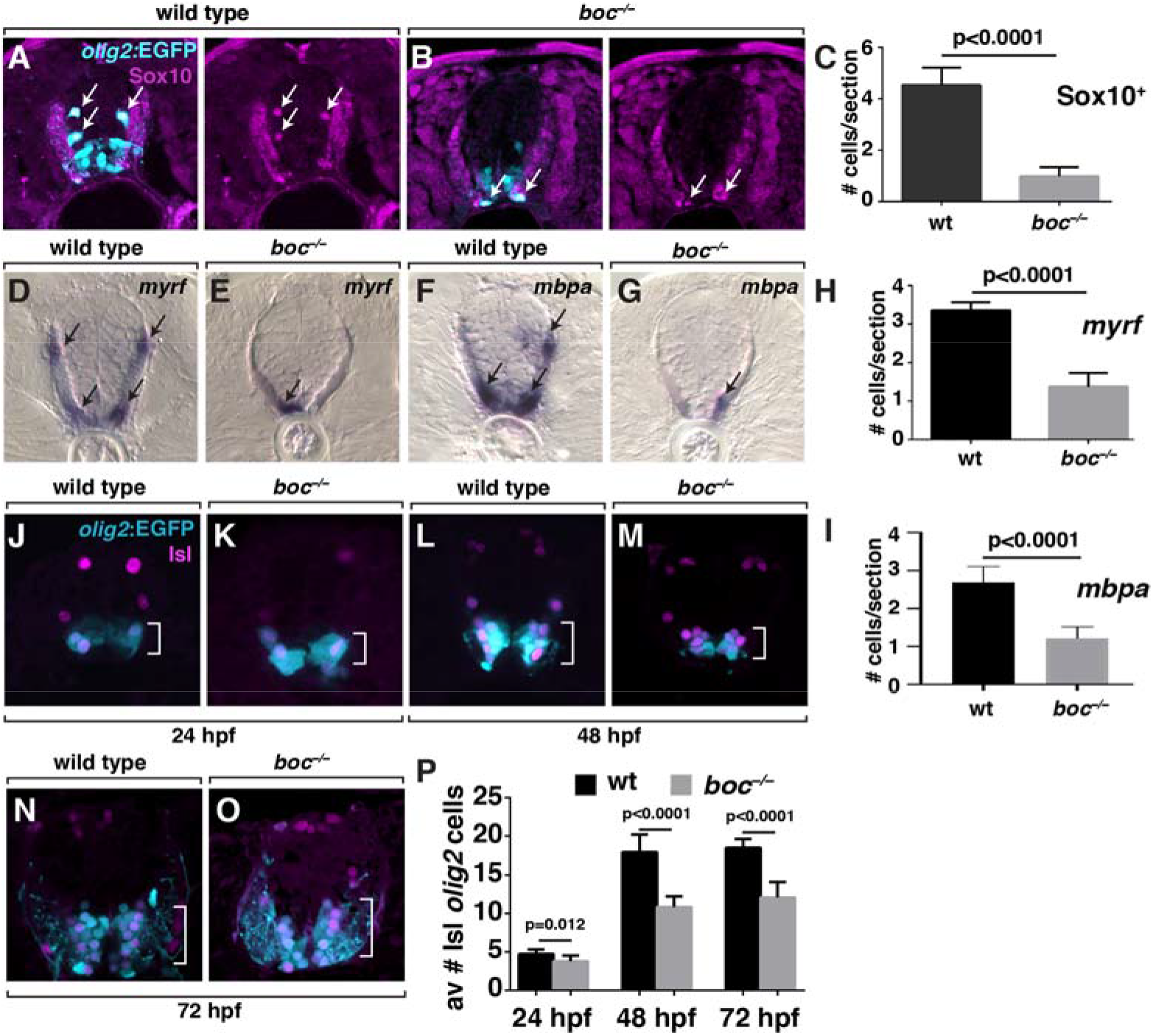
*boc* mutant animals have deficits of oligodendrocyte lineage cells and motor neurons. All panel images show transverse sections of trunk spinal cord oriented with dorsal to the top. (A,B) *olig2*:EGFP reporter expression (cyan) combined with immunohistochemistry to detect Sox10 (magenta) in 3 dpf wild-type and *boc* mutant larvae. Arrows mark double labeled oligodendrocyte lineage cells. (C) *boc* mutant larvae have fewer Sox10^+^ oligodendrocyte lineage cells than wild type. (D-G) 3 dpf wild-type and *boc* mutant larvae processed to detect *myrf* and *mbpa* transcripts as markers of pre-myelinating oligodendrocytes and myelinating oligodendrocytes, respectively. (H) *boc* mutant larvae have fewer *myrf*^+^ cells than wild type. (I) *boc* mutant larvae have a deficit of *mbpa*^+^ oligodendrocytes relative to wild type. (J-O) *olig2*:EGFP reporter expression (cyan) combined with immunohistochemistry to detect Isl (magenta) in wild-type and *boc* mutant embryos and larvae. Double labeled cells are motor neurons (bracketed areas). (P) *boc* mutant embryos have normal number of motor neurons relative to wild type at 24 hpf, but a deficit at 48 and 72 hpf. Data are presented as mean ± SEM and evaluated using two-tailed, unpaired *t*-test with a 95% confidence interval. n = 10 embryos for each genotype and condition.

Oligodendrocyte lineage cells arise from *olig2*^*+*^ pMN progenitors following formation of motor neurons. To determine if loss of *boc* function also affects motor neuron production we performed immunohistochemistry to detect Isl protein. In combination with *olig2:*EGFP expression, anti-Isl labeling provides a definitive marker of motor neurons. At 24 hpf, during the peak of motor neuron production, the numbers of spinal motor neurons in *boc* mutant embryos and their wild-type siblings were similar (Figure 2I,J,O). At 48 hpf, following completion of embryonic motor neuron formation, *boc* mutant embryos had a substantial deficit of motor neurons compared to wild-type embryos (Figure 2K,L,O). This deficit persisted through 3 dpf (Figure 2M-O). Altogether, these data indicate that *boc* function is necessary to produce normal numbers of motor neurons and oligodendrocytes.

### boc function is required to maintain pMN progenitor cell identity

One possible explanation for the data above is that *boc* function is necessary to maintain progenitors that produce motor neurons and oligodendrocyte lineage cells. To investigate this possibility, we performed in situ RNA hybridization to detect expression of *nkx2.2a*, which marks p3 progenitors, the ventral-most progenitor domain of the spinal cord, and *olig2*, which marks pMN progenitors immediately dorsal to p3 cells. At 24 hpf, wild-type and *boc* mutant larvae appeared to express both *nkx2.2a* and *olig2* similarly (Figure 3A-D), suggesting that ventral spinal cord patterning and ventral progenitor domain formation are not affected by loss of *boc* function. At 48 hpf, *nkx2.2a* expression again appeared to be similar in wild-type and *boc* mutant embryos (Figure 3E,F). However, whereas pMN cells of wild-type embryos continued to express *olig2* (Figure 3G)*, boc* mutant embryos were devoid of *olig2* expression (Figure 3H). At 72 hpf, *boc* mutant larvae continued to express *nkx2.2a* equivalently to wild type (Figure 3I,J) but did not express *olig2* at detectable levels, compared to the low levels of message still evident in wild-type larvae (Figure 3K,L). We interpret these data to mean that Boc is not necessary to pattern the ventral spinal cord but instead is required to maintain *olig2* expression through the period of neurogenesis and gliogenesis.

**Figure 3.**
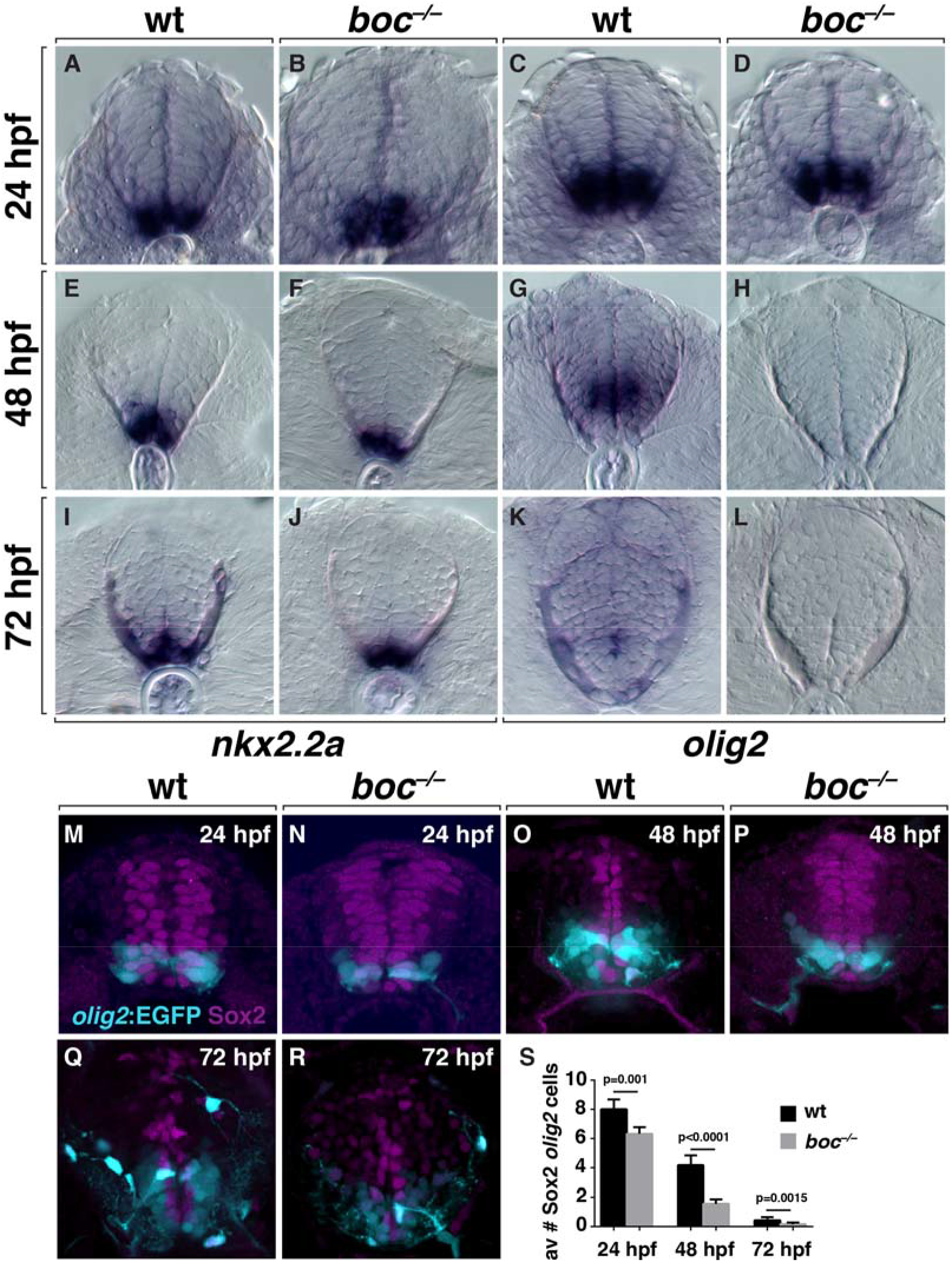
*boc* function maintains pMN progenitors. All image panels show transverse trunk spinal cord sections oriented with dorsal to the top. (A-L) Wild-type and *boc* mutant embryos and larvae processed to detect *nkx2.2a* and *olig2* mRNA expression as markers of p3 and pMN progenitors, respectively. (A-D) At 24 hpf, wild-type and mutant embryos express *nkx2.2a* and *olig2* expression similarly. (E-F) At 48 hpf, mutant embryos express *nkx2.2a* similarly to wild type but do not express *olig2*, in contrast to wild-type embryos. (I-L) At 72 hpf, wild-type and mutant larvae continue to express *nkx2.2a* similarly but express little *olig2* mRNA. (M-R) *olig2*:EGFP reporter expression (cyan) in combination with immunohistochemistry to detect Sox2 (magenta) as a marker of progenitors. (S) *boc* mutant embryos and larvae have fewer *olig2*:EGFP^+^ Sox2^+^ pMN progenitors at all stages. Data are presented as mean ± SEM and evaluated using two-tailed, unpaired *t*-test. n = 10 embryos for each genotype and condition.

Is *boc* function required to maintain the population of spinal cord progenitors, or does it more specifically maintain the identity of *olig2*^+^ pMN progenitors? To address this question we used immunohistochemistry to detect expression of Sox2, which marks spinal cord progenitors, in combination with *olig2*:EGFP. The number and distribution of Sox2^+^ cells appeared to be similar between genotypes at 24, 48 and 72 hpf (Figure 3M- R). However, *boc* mutant embryos and larvae had a deficit of Sox2^+^ *olig2:*EGFP^+^ progenitors at each stage (Figure 3M-S). We note that this assay probably overestimates the number of Sox2^+^ *olig2:*EGFP^+^ cells at 48 and 72 hpf because of the stability of the EGFP signal relative to RNA detection (compare Figure 3H,L,P,R). These data indicate that *boc* function is required to maintain pMN progenitor identity, rather than to maintain a neural progenitor state.

### boc modulates Shh signaling across the spinal cord dorsoventral axis

Our previous data have shown that sustained Shh signaling is required to maintain *olig2*^*+*^ pMN progenitors and promote OPC formation (Park *et al.* 2004; Ravanelli and Appel 2015). Therefore, the premature loss of pMN progenitors by *boc* mutant embryos might result from abnormal Shh signaling. As a first test of this possibility, we examined expression of *shha*. At 24 and 72 hpf, wild-type and *boc* mutant embryos expressed *shha* mRNA similarly (Figure 4A-D) indicating that loss of *boc* function does not affect expression of the Shh ligand at the level of transcription. Next, we tested expression of a *gli1* reporter gene, *8xGliBS:mCherry-NLS,* which serves as an indicator of high levels of Shh signaling activity in the ventral spinal cord (Mich *et al.* 2014). At 24 hpf, cells that expressed the highest levels of *8xGliBS:*mCherry-NLS fluorescent protein in wild-type embryos were ventral to *olig2*:EGFP^+^ pMN cells, whereas numerous pMN cells expressed *8xGliBS:*mCherry-NLS at lower levels (Figure 4E,E’). By contrast, cells that expressed *8xGliBS:*mCherry-NLS at high levels were evident ventral to pMN cells in 24 hpf *boc* mutant embryos but little *gli1* reporter expression was evident with pMN cells (Figure 4F,F’). At 48 hpf, *boc* mutant embryos similarly expressed lower levels of *8xGliBS:*mCherry-NLS than wild-type embryos (Figure 4G-H’’). Together these data suggest that *boc* function promotes and sustains Shh signaling in pMN cells.

**Figure 4.**
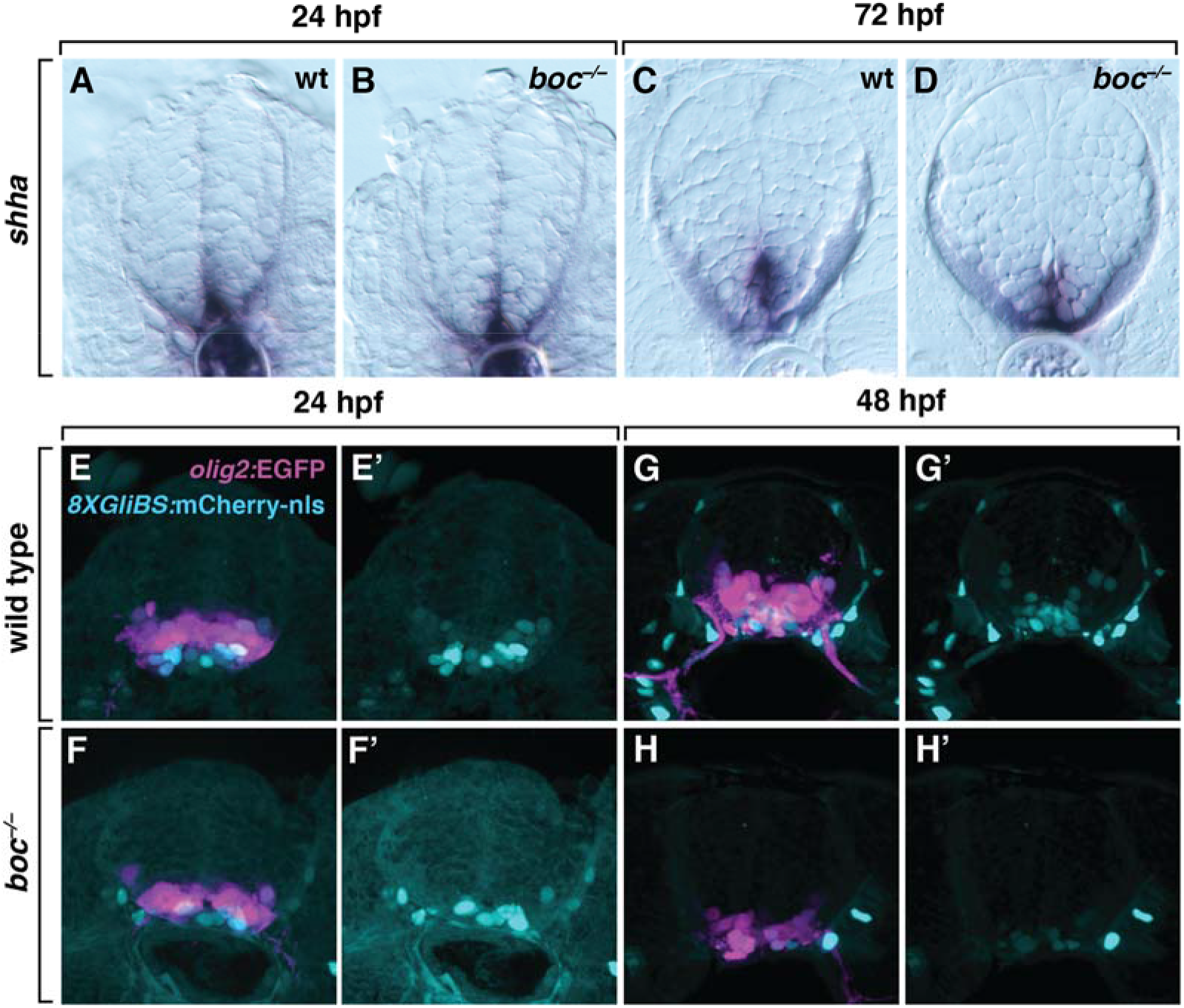
*boc* promotes Shh signaling response in the ventral spinal cord. All panels show transverse sections at the level of the trunk spinal cord. (A-D) Representative images of wild-type (wt) and *boc* mutant spinal cords processed to detect *shha* transcripts (blue) using in situ RNA hybridization. Transcript amount and distribution appear similar between wild-type and mutant spinal cords at 24 and 72 hpf. (E-H’) Representative images of wild-type and *boc* mutant spinal cords showing *olig2*:EGFP (magenta) and *8xGliBS*:mCherry-nls (cyan) expression. Expression of the *gli1* reporter appears reduced in mutant embryos relative to wild-type embryos.

Previous data from mouse embryos indicated that dorsal spinal cord cells express *Boc* at high level, with less expression evident in the ventral spinal cord (Mulieri *et al.* 2002; Tenzen *et al.* 2006; Allen *et al.* 2011). To examine zebrafish *boc* expression in relationship to Shh signaling activity and pMN cells, we used a highly sensitive fluorescent detection method for in situ RNA hybridization to detect *boc* mRNA in combination with *olig2* and *ptch2*, which provides a readout of Shh signaling. At 30 hpf *boc* mRNA was evident as a dorsal to ventral gradient, with the highest levels of expression in the dorsal spinal cord (Figure 5A,A”). *ptch2* mRNA was expressed in a reciprocal ventral to dorsal gradient (Figure 5A,A’). Notably, pMN progenitors, marked by *olig2* expression, expressed *ptch2* as well as a low level of *boc* transcripts (Figure 5A). By 48 hpf, spinal cord distribution of *boc* mRNA was evident as a dorsoventral stripe corresponding to the position progenitor cells, including pMN progenitors (Figure 5B,B”) and overlapping with *ptch2* expression (Figure 5B,B’). By 72 hpf, little *olig2* expression remained, indicative of pMN progenitor depletion, and *ptch2* and *boc* expression were diminished (Figure 5C-C”). These data are consistent with the possibility that *boc* functions within ventral spinal cord to maintain pMN progenitors and specify OPC fate.

**Figure 5.**
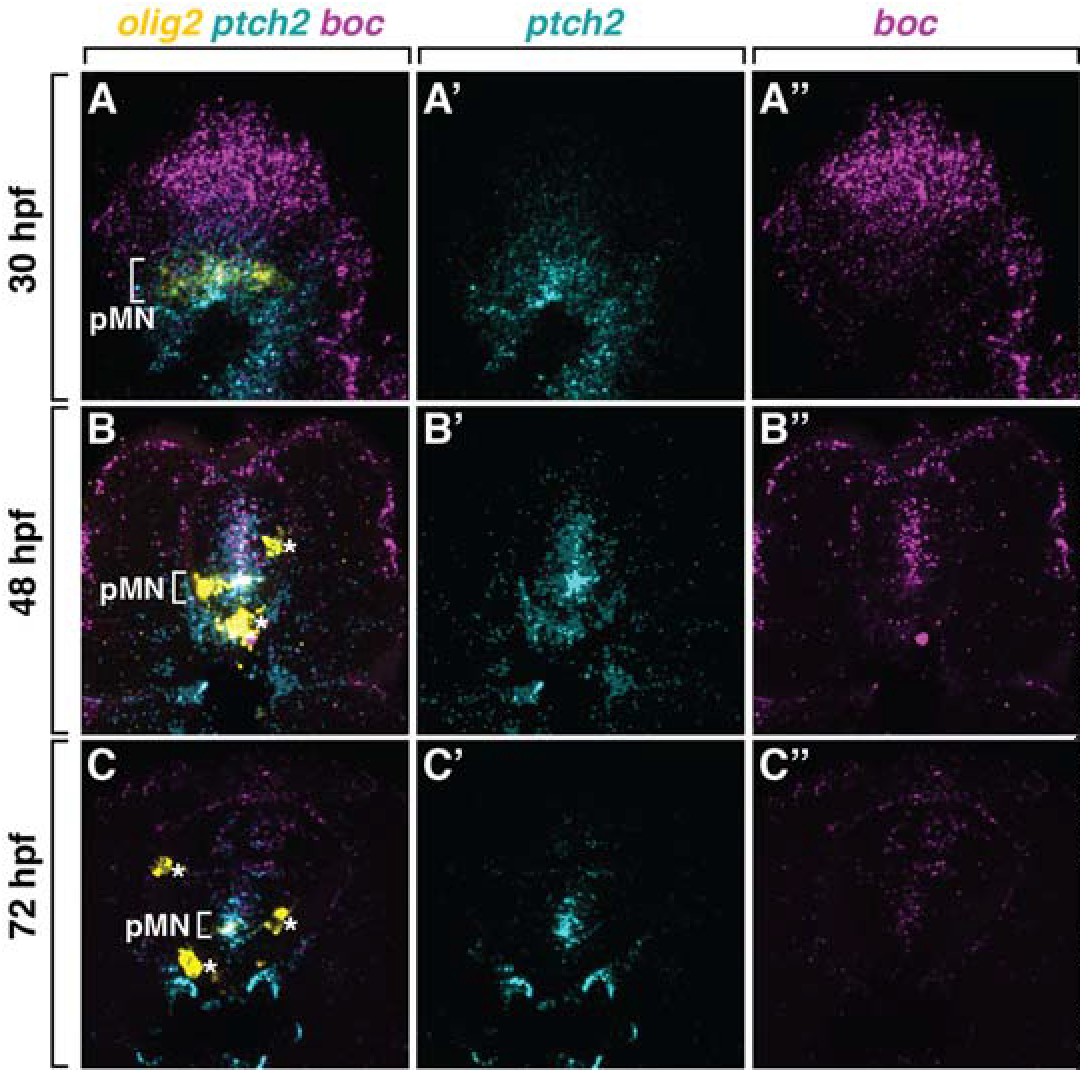
Spinal cord cells express *boc* in a ventral to dorsal gradient. All panels show transverse sections of wild-type embryos and larvae through the trunk spinal cord, with dorsal up, processed to detect *olig2* (yellow), *ptch2* (cyan) and *boc* (magenta) mRNA. At 30 and 48 hpf, pMN cells (bracket) express *ptch2* and low levels of *boc.*

To learn how Boc function regulates Shh signaling within the spinal cord we examined expression of *ptch2* in combination with *boc* and *olig2* in *boc* mutant embryos using fluorescent in situ RNA hybridization. At all stages *boc* mRNA expression appeared to be similar in *boc* mutant embryos (Figure 6) and wild-type embryos at comparable stages (Figure 5). *olig2* expression appeared to be normal at 30 hpf (Figure 6A) but depleted at 48 and 72 hpf (Figure 6B,C), consistent with our observations above (Figure 3D,H,L). Remarkably, at 30 hpf, *ptch2* expression appeared to be expanded more dorsally in mutant embryos (Figure 6A,A’), compared to wild type (Figure 5A,A’). At 48 and 72 hpf *ptch2* expression appeared to be more uniform in the spinal cords of *boc* mutant embryos (Figure 6B,C) compared to the graded distribution evident in wild-type embryos (Figure 5B,C). To validate our visual observations, we quantified fluorescence intensities across the dorsoventral spinal cord axis of 30 hpf wild-type and *boc* mutant embryos. These data show that peak *olig2* and *ptch2* expression coincide at approximately 20 µm from the ventral limit of the spinal cord in wild-type embryos (Figure 6D,F). *olig2* expression was similar in wild-type and mutant embryos (Figure 6D), confirming that loss of *boc* function does not impair formation of the pMN domain. By contrast, the graded distribution of *boc* RNA was flattened in *boc* mutant embryos, with higher levels ventrally and lower levels dorsally that in wild-type embryos (Figure 6E). Finally, the ventral to dorsal gradient of *ptch2* expression was shifted dorsally in *boc* mutant embryos relative to wild-type embryos (Figure 6F). These observations suggest that Boc function helps refine precisely graded Shh signaling within the ventral spinal cord to maintain pMN progenitors and produce late-born motor neurons and oligodendroctyes.

**Figure 6.**
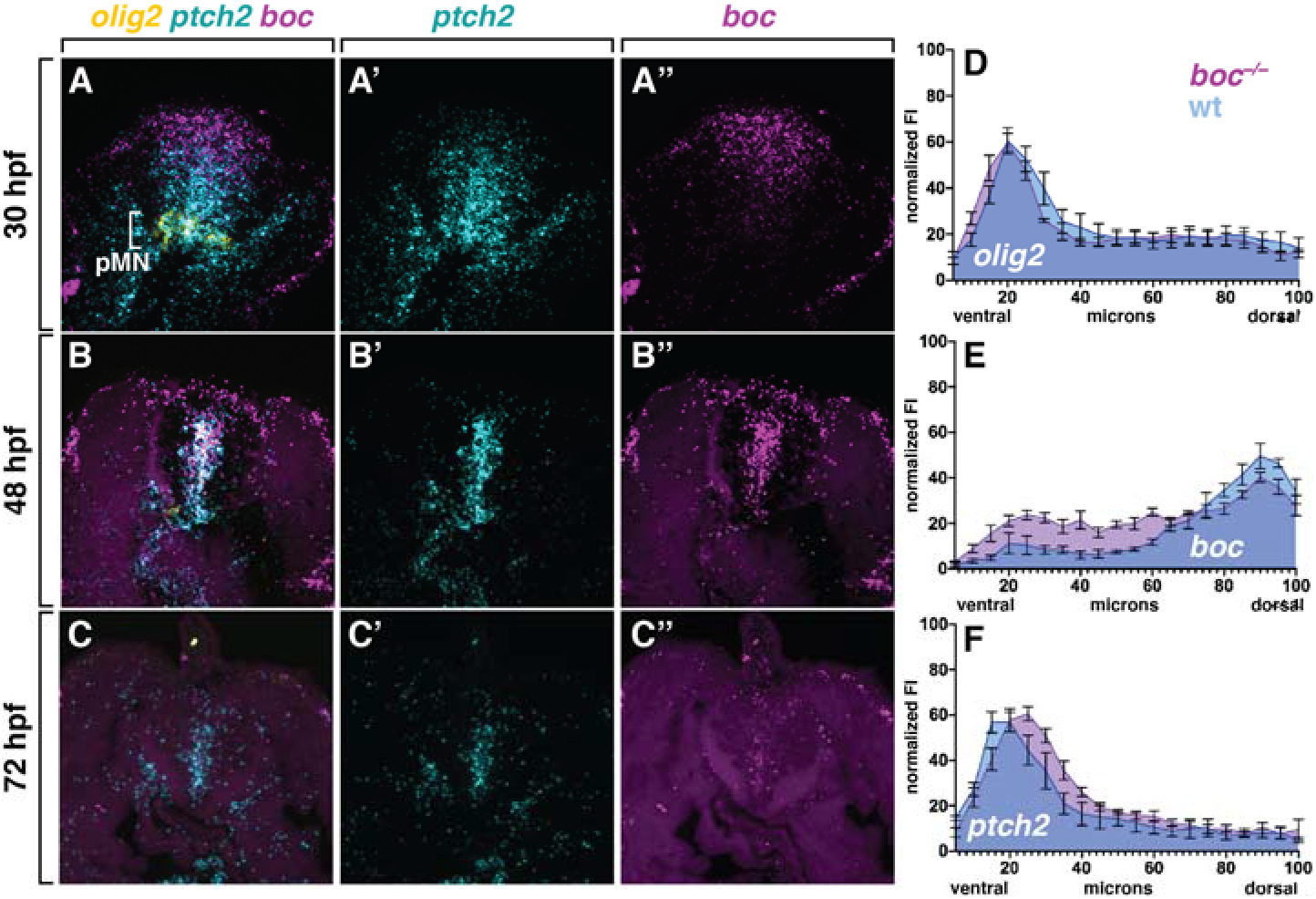
Loss of *boc* function alters the spinal cord Shh signaling gradient. (A-C) All panels show transverse sections of *boc* mutant embryos and larvae through the trunk spinal cord, with dorsal up, processed to detect *olig2* (yellow), *ptch2* (cyan) and *boc* (magenta) mRNA. At 30 and 48 hpf, expression appears to extend more dorsally and to be less graded across the ventral to dorsal axis than wild type (compare to Figure 5). (D-F) Normalized fluorescence intensity across the spinal cord dorsoventral axis for wild-type (light blue) and *boc* mutant (magenta) embryos at 30 hpf. (D) The peak of *olig2* RNA expression is similar in wild-type and mutant embryos. (E) *boc* RNA expression is higher ventrally and lower dorsally in *boc* mutant embryos compared to wild type. (F) The peak of *ptch2* expression is shifted dorsally in *boc* mutant embryos relative to wild type. The y axis fo each graph shows normalized fluorescence intensity (FI) and the x axis represents distance in from the ventral to the dorsal limits of the spinal cord. Data are presented as ± SEM. n = 5 embryos.

## Discussion

In the developing ventral spinal cord, Shh signaling patterns the dorsoventral axis to specify discrete subpopulations of neural progenitors (Sagner & Briscoe, 2019) and subsequently causes pMN progenitors to switch from producing motor neurons to OPCs (Danesin and Soula 2017). By investigating a mutant allele of *boc* uncovered in a forward genetic screen we have learned that Boc helps maintain Shh signaling at levels necessary to preserve pMN progenitors for OPC formation by shaping the Shh signaling gradient.

One of the main findings of our work is that a missense mutation within the FNIII(3) domain of Boc produces a loss-of-function phenotype similar to that resulting from the *boc*^*ty54*^ allele, which is predicted to cause a truncation of the Boc protein that eliminates two of four immunoglobulin domains and all three Fn type-III domains (Bergeron *et al.* 2011). A previous study provided biochemical evidence that Shh binds to the Boc FNIII(3) domain and used a gain-of-function approach in chick neural tube to show that the FNIII(3) domain necessary for the ability of Boc to enhance Shh signaling (Tenzen *et al.* 2006). Thus, our work helps confirm the conclusion that the FNIIII(3) domain is the principal site of functional interaction between Boc and Shh.

Our analysis of the *boc* mutant phenotype revealed that homozygous mutant larvae have a deficit of oligodendrocyte lineage cells. Mutant larvae had deficits of both total oligodendrocyte lineage cells and differentiating oligodendrocytes, suggesting that fewer OPCs are formed by neural progenitors in the absence of *boc* function, resulting in formation of fewer myelinating cells. Similarly, P0-P1 *Boc* mutant mouse pups had a deficit of cells expressing the OPC markers Olig2 and PDGFRa in the dorsal forebrain (Zakaria *et al.* 2019). Thus, progenitor populations in different regions of the developing nervous system may have a common requirement for Boc function in formation of OPCs from neural progenitors. The developing dorsal forebrain of *Boc* mutant mice also had fewer differentiating oligodendrocytes and less myelin than wild-type mice, although the ratio of oligodendrocytes to OPCs was similar to that of wild type (Zakaria *et al.* 2019). Thus, in both zebrafish and mice Boc function appears to be required for OPC production and not for oligodendrocyte differentiation.

Why do *boc* mutant zebrafish produce fewer spinal cord OPCs? Another main finding of our work is that Boc is required to maintain pMN progenitors as they transition from producing motor neurons to OPCs. Specifically, we found that *boc* mutant embryos initially expressed *olig2* and *nkx2.2a* normally, indicating that ventral spinal cord patterning does not require Boc function, but that *olig2* transcripts were prematurely depleted from mutant spinal cords relative to wild type. Previously, we showed that treating 24 or 30 hpf zebrafish embryos with cyclopamine, a pharmacological inhibitor of Shh signaling, similarly caused premature depletion of *olig2*-expressing cells and a deficit of OPCs (Park *et al.* 2004; Ravanelli and Appel 2015). Taken together, these results suggest that Boc maintains Shh signaling activity within ventral spinal cord cells to preserve a population of *olig2*^+^ pMN progenitors for OPC formation.

Our interpretation that Boc maintains Shh signaling in the ventral spinal cord was confounded by data showing that *Boc* expression is evident in dorsal but not ventral spinal cord of mouse embryos (Mulieri *et al.* 2002; Tenzen *et al.* 2006; Allen *et al.* 2011). By using a highly sensitive fluorescent detection method for in situ RNA hybridization we found that, in fact, zebrafish pMN progenitors express low levels of *boc* transcripts. Additionally, our in situ RNA hybridization data revealed that *boc* transcripts are expressed in a graded, dorsal-high to ventral-low pattern that is inverse to the pattern of *ptch2* expression, an indicator of Shh signaling activity. Because Boc promotes Shh signaling and Shh signaling negatively regulates *Boc* expression (Tenzen *et al.* 2006), our results now suggest that a cross-regulatory relationship between Shh signaling and Boc helps tunes the Shh signaling gradient in the spinal cord.

The premature loss of *olig2*^+^ pMN cells from *boc* mutant embryos led us to predict that Boc maintains Shh signaling within pMN progenitors. Consistent with this, *boc* mutant embryos expressed a *gli1* reporter gene at reduced levels. However, our in situ RNA hybridization experiments produced a surprising result. Specifically, *boc* mutant embryos expressed high levels of *ptch2*, indicating that Shh signaling was preserved or even enhanced. Consistent with this interpretation, the level of *boc* mRNA expression, which is negatively regulated by Shh signaling, appeared to be reduced. Importantly, quantification of expression levels across the dorsoventral axis showed that the peak of *ptch2* expression was shifted dorsally with respect to *olig2* expression at 30 hpf, prior to OPC specification. One possible interpretation for this observation is that even low levels of Boc expressed by pMN cells limits the dorsal spread of Shh, helping concentrate Shh signaling within the pMN domain. In the absence of Boc, Shh might be free to move more dorsally, shifting the signaling gradient. Consequently, the level of Shh signaling within the pMN domain might be reduced below a threshold necessary to specify formation of OPCs.

The main limitation of our study is that we do not know the fate of progenitors in *boc* mutant embryos that normally would develop as OPCs. We previously showed that cells that produce OPCs are recruited from progenitors dorsal to the pMN domain by sustained Shh signaling (Ravanelli and Appel 2015). One possibility is that in the absence of *boc* function these progenitors instead develop as interneurons or glia characteristic of the p2 progenitor domain, which lies just dorsal to the pMN domain. This can be addressed in future work using cell fat mapping approaches.

## Materials and Methods

### Zebrafish lines and husbandry

All animal work was approved by the Institutional Animal Care and Use Committee (IAUCUC) at the University of Colorado School of Medicine. All non-transgenic embryos were obtained from pairwise crosses of males and females from the AB strain. Embryos were raised at 28.5°C in E3 media (5 mM NaCl, 0.17 mM KCl, 0.33 mM CaCl_2_, 0.33 mM MgSO_4_ at pH 7.4, with sodium bicarbonate), sorted for good health and staged accordingly to developmental morphological features and hours post-fertilization (hpf) (Kimmel *et al.* 1995). Developmental stages are described in the results section for individual experiments. Sex cannot be determined at embryonic and larval stages. The transgenic lines used were *Tg(olig2:EGFP)*^*vu12*^ (Shin et al., 2003), and *Tg(8xGliBS:mCherry-NLS)*^*st1001Tg*^ (Mich *et al.* 2014). All transgenic embryos were obtained from pairwise crosses of males females from the AB strain to males or females of each transgenic line used.

### boc mutant allele identification

We created a mapping cross by mating *co25*^*+/−*^ fish, which were from the AB strain, to WIK strain fish and raising the progeny to adulthood. We then mated pairs of heterozygous map cross adults and collected 24 wild-type larvae and 24 mutant larvae at 5 dpf, from which we prepared and pooled genomic DNA. Next, we performed bulk segregant analysis (Knapik *et al.* 1998) using 223 simple sequence-length polymorphism markers. This placed the *co25* locus between 36 cM (Z23011) and 53.4 cM (Z13229) on chromosome 24. We selected *boc*, located at 44.1 cM, as a candidate gene because of the similarity between homozygous *boc* mutant embryos (Bergeron *et al.* 2011) and *co25* mutant embryos. We prepared pooled total RNA from 10 homozygous mutant larvae and 10 wild-type larvae at 4 dpf, synthesized cDNA and sequenced PCR products amplified from overlapping fragments of the *boc* coding region.

### boc genotyping

The *co25* allele introduces a SfcI restriction site. For genotyping, we amplified a 97 bp fragment using the PCR primers 5’-AACACCCCCATTTATGGATTC-3’ and 5’-AACAAGACTGACCTTCCACCA-3’. The fragments were digested with SfcI and run on 3% agarose gels. Homozygous wild type animals produced a single 97 bp band, heterozygotes produced 97, 77 and 20 bp bands and homozygous mutants produced 77 and 20 bp bands.

### Fluorescent In situ RNA Hybridization

Fluorescent in situ RNA hybridization was performed using the RNAScope Multiplex Fluorescent V2 Assay Kit (Advanced Cell Diagnostics; ACD) on 12 μm thick paraformaldehyde-fixed and agarose embedded cryosections according to manufacturer’s instructions with the following modification: slides were covered with parafilm for all 40°C incubations to maintain moisture and disperse reagents across the sections. The zebrafish *olig2* C1, *ptch2*-C2, and *boc1*-C3 transcript probes were designed and synthesized by the manufacturer and used at 1:50 dilutions. Transcripts were fluorescently labeled with Opal520 (1:1500), Opal570 (1:500) and Opal650 (1:1500) using the Opal 7 Kit (NEL797001KT; Perkin Elmer).

### Chromogenic In situ RNA Hybridization

Chromogenic in situ RNA hybridizations were performed as described previously (Hauptmann and Gerster 2000). Probes included *nkx2.2a* (Barth and Wilson 1995)*, olig2* (Park *et al.* 2002), *shha* (Krauss *et al.* 1993) *myrf* (Scott *et al.* 2020) and *mbpa* (Brösamle and Halpern 2002). Plasmids were linearized with appropriate restriction enzymes and cRNA preparation was carried out using Roche DIG-labeling reagents and T3, T7 or SP6 RNA polymerases (New England Biolabs). After staining, embryos were embedded in 1.5% agar/5% sucrose and frozen over dry ice. 20 μm transverse sections were cut using the Leica CM 1950 cryostat (Leica Microsystems), collected on microscope slides and mounted with 75% glycerol.

### Immunohistochemistry

Larvae were fixed using 4% paraformaldehyde/1X PBS overnight at 4°C. Embryos were washed with 1X PBS, rocking at room temperature and embedded in 1.5% agar/5% sucrose, frozen over dry ice and sectioned in 20 or 15 µm transverse increments using a cryostat microtome. Slides were place in Sequenza racks (Thermo Scientific), washed 3×5 min in 0.1%Triton-X 100/1X PBS (PBSTx), blocked 1 hour in 2% goat serum/2% bovine serum albumin/PBSTx and then placed in primary antibody (in block) overnight at 4°C. The primary antibodies used include: rabbit anti-Sox10 (1:500) (Park *et al.* 2005), rabbit anti-Sox2 (1:500, # ab997959, Abcam) and mouse anti-Islet (1:500; Developmental Studies Hybridoma Bank, AB2314683). Sections were washed for 1 hours at room temperature with PBSTx and then incubated for 2 hr at room temperature with secondary antibodies at a 1:200 dilution in block. The secondary antibodies used include: AlexaFluor 568 anti-rabbit (Invitrogen A11011) and AlexaFluor 568 anti-mouse (Invitrogen A11004). Sections were washed for 1 hr with PBSTx and mounted in VectaShield (Vector Laboratories).

### Imaging

Fixed sections of embryos and larvae were imaged on a Zeiss CellObserver SD 25 spinning disk confocal system (Carl Zeiss). Cell counts were collected using a 20x objective (n.a. 0.8) and representative images were collected using a 40x oil immersion objective (n.a. 1.3). Chromogenic RNA in situ hybridization sections were imaged using differential interference contrast optics and a Zeiss AxioObserver compound microscope (Carl Zeiss). Cell counts and representative images were acquired at 40X (n.a. 0.75). Images are reported as extended z-projections or a single plane (in situ RNA hybridization) collected using Zen (Carl Zeiss) imaging software. Image brightness and contrast were adjusted in Photoshop (Adobe) or ImageJ (National Institutes of Health).

### Quantification of fluorescent images

Quantification of fluorescent RNA in situ hybridization images was carried out using z-projections collected at identical exposures using a confocal microscope. Using Fiji (Schindelin *et al.* 2012), ten 0.5 um z intervals were Z-projected using the “Sum of Slices” function. A 20-point line was then drawn across the center of the spinal cord from the ventral floor to the dorsal roof. Average gray values across the spinal cord were measured for *olig2*, *ptch2* and *boc* by recording the data stored in histograms for each channel. All gray values were normalized for each channel within each analyzed section on a 0-100% scale using the following function:

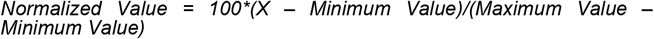

The distance for each analyzed section was scaled, with 0 representing the most ventral measurement and 100 designating the most dorsal measurement. After normalizing all collected data, the average normalized fluorescence intensity was calculated across the spinal cord at 5% distance intervals.

### Data Quantification and Statistical Analysis

We plotted all data and performed all the statistical analyses in GraphPad Prism. All data are expressed as mean±SEM. For statistical analysis, we used an unpaired Student’s two-tailed *t*-test with a 95% confidence interval. Unless otherwise stated, all graphs represent data collected from one experimental replicate, sampling fish from multiple cross with no inclusion or exclusion criteria.

### Data and Reagent Availability

All data, reagents and zebrafish lines are available by communication with the corresponding author.

## End Matter

### Author Contributions and Notes

M.W., A.M.R. and B.A. designed research, C.A.K, M.W., A.M.R., K.S., M.R.A. and B.A. performed research, C.K. and K.S. analyzed data and B.A. wrote the paper.

The authors declare no conflict of interest.

## Acknowledgments

We thank members of the Appel lab for discussions. This work was supported by a National Institutes of Health grant (NS406668) and a gift from the Gates Frontiers Fund to B.A. A.M.R. was supported by the National Institutes of Health (National Cancer Institute; T32 CA08208613). K.S. was supported by the National Institutes of Health (F31 NS116922). M.R.A was supported by a Children’s Hospital Colorado summer research internship. The University of Colorado Anschutz Medical Campus Zebrafish Core Facility was supported by a National Institutes of Health grant (P30 NS048154). The anti-Isl antibody, developed by T.M. Jessell and S. Brenner-Morton, was obtained from the Developmental Studies Hybridoma Bank, created by the NICHD of the NIH and maintained at The University of Iowa, Department of Biology, Iowa City, IA 52242.

